# Axonal PLC-γ activity is required for BDNF long-distance signaling

**DOI:** 10.1101/2023.11.10.566602

**Authors:** G Moya-Alvarado, X Valero-Peña, A Aguirre-Soto, FC Bronfman

## Abstract

Brain-derived neurotrophic factor (BDNF) and its receptor tyrosine kinase receptor B (TrkB) are important signaling proteins that regulate dendritic growth and maintenance in the central nervous system (CNS). After binding of BDNF, TrkB is endocytosed into endosomes and continues signaling within the cell soma, dendrites, and axon. We showed that BDNF signaling initiated in axons triggers long-distance signaling, inducing dendritic arborization in a CREB-dependent manner in cell bodies, processes that depend on axonal dynein and TrkB activities. The binding of BDNF to TrkB triggers the activation of different signaling pathways, including the ERK1/2, PLC-γ and PI3K-mTOR pathways, to induce dendritic growth and synaptic plasticity. How TrkB downstream pathways regulate long-distance signaling is unclear. Here, we studied the role of PLC-γ-Ca^2+^ in BDNF/TrkB-induced long-distance signaling using compartmentalized microfluidic cultures. We found that dendritic branching and CREB phosphorylation induced by axonal BDNF stimulation require the activation of PLC-γ in the axons of cortical neurons. Locally on axons, BDNF increases PLC-γ phosphorylation and induces intracellular Ca^2+^ waves in a PLC-γ-dependent manner. Moreover, the transport of BDNF-containing signaling endosomes to the cell body was dependent on PLC-γ activity and intracellular Ca^2+^ stores. Furthermore, the activity of PLC-γ is required for BDNF-dependent TrkB endocytosis, suggesting a role for the TrkB/PLC-γ signaling pathway in axonal signaling endosome generation.

## INTRODUCTION

Brain-derived neurotrophic factor (BDNF) and its receptor TrkB are major regulators of dendritic branching in cortical and hippocampal neurons (Cheung et al., 2007, Horch and Katz, 2002, Xu et al., 2000). TrkB is located both in the somatodendritic compartment and axonal compartment (AC) of neurons (Arimura et al., 2009), and BDNF is released in both dendrites and axons (Matsuda et al., 2009). Upon binding to TrkB, BDNF activates three main downstream signaling molecules, phospholipase C- γ (PLC-γ), mitogen-activated protein kinases (MAPKs) and phosphoinositide 3-kinase (PI3K) (Reichardt, 2006), to regulate dendritic and axonal growth (Huang and Reichardt, 2003, Gonzalez et al., 2016), synaptic transmission (Kang and Schuman, 1995, Zhang et al., 2013) and learning and memory (Alonso et al., 2002b, Alonso et al., 2002a, Egan et al., 2003). Extracellular application of BDNF induces an increase in cytoplasmic Ca^2+^ in a PLC-γ-dependent manner (Li et al., 2005). Furthermore, it has been shown that Ca^2+^ influx increases dendritic arborization via activation of the transcriptional program that involves activation of calmodulin kinase IV (CAMKIV) and phosphorylation of the cAMP response element-binding protein (CREB) transcription factor (Redmond et al., 2002). BDNF promotes the release of intracellular Ca^2+^ stores by activation of the inositol 3,4,5-triphosphate receptor (IP3R) and the entry of extracellular Ca^2+^ by opening transient receptor-potential cation channel subfamily C (TRPC) (Li et al., 2005, Leal et al., 2015). Moreover, BDNF-dependent long-term potentiation in the hippocampus requires both pre- and post-synaptic PLC-γ activity (Gärtner et al., 2006, Gruart et al., 2007).

In cortical and hippocampal neurons, the binding of BDNF to TrkB promotes its dimerization and autophosphorylation, inducing the internalization of the receptor into signaling endosomes (Cosker and Segal, 2014, Scott-Solomon and Kuruvilla, 2018, Bronfman et al., 2014, Moya-Alvarado et al., 2022). We have previously shown that BDNF axonal signaling promotes dendritic arborization in a TrkB-dependent manner (Moya-Alvarado et al., 2023). TrkB is endocytosed in axons and retrogradely transported to the cell body by the molecular motor dynein. In the cell body, activated TrkB upregulates CREB and PI3K/mTOR activity, increasing the transcription and translation of proteins (Moya-Alvarado et al., 2023). Although long-distance signaling of BDNF has been described in cortical neurons (Cohen et al., 2011, Zhou et al., 2012), little is known about the cellular mechanism and the downstream signaling pathways regulating this process. Several lines of evidence support the participation of the PLC-γ and increased intracellular Ca^2+^ in long-distance signaling mediated by BDNF. It has been reported that in hippocampal neurons, the internalization of the BDNF/TrkB complex is regulated by an increase in Ca^2+^ influx through N-methyl-d-aspartate receptors (NMDARs) (Du et al., 2003). Furthermore, the axonal stimulation of retinal ganglionic cells (RGCs) with BDNF promotes a retrograde potentiation of retinal synapses in a TrkB- and PLC-γ-dependent manner (Du and Poo, 2004, Lom et al., 2002a), suggesting that PLC-γ can participate in BDNF long-distance signaling. Interestingly, in the axons of sympathetic neurons, TrkA via PLC-γ regulates the activation of calcineurin to dephosphorylate dynamin 1, coordinating receptor endocytosis and axonal growth (Bodmer et al., 2011). Taken together, these results suggest that activation of the PLC-γ/Ca^2+^ pathway participates in the endocytosis and trafficking of Trks receptors.

Here, we show that BDNF long-distance signaling depends on the axonal activity of PLC-γ. Using compartmentalized cultures of cortical neurons, we found that axonal PLC-γ activity in axons is required for dendritic arborization and CREB phosphorylation in cell bodies. Consistent with other works, our work reveals that BDNF increases axonal Ca^2+^ levels in a PLC-γ-dependent manner (Du and Poo, 2004). Moreover, BDNF increases the activation of PLC-γ in axons, and the transport of signaling endosomes from axons to cell bodies was dependent on both PLC-γ activity and intracellular calcium. Furthermore, we show that the PLC-γ pathway is required for TrkB endocytosis, suggesting that axonal PLC-γ can regulate the generation of signaling endosomes for retrograde BDNF/TrkB signaling.

## RESULTS

### Axonal PLC-γ regulates BDNF/TrkB long-distance signaling in cortical neurons

Previously, we reported that axonal stimulation with BDNF promotes dendritic arborization in a CREB-dependent manner. Interestingly, when characterizing the downstream TrkB signaling pathways, we found that PI3K activity is required in the cell body but not in the axons of cortical neurons to induce dendritic arborization and mTOR activation in cell bodies (Moya-Alvarado et al., 2023). One downstream pathway involved in TrkB signaling in axons is PLC-γ, as shown for RGCs and sympathetic neurons (Bodmer et al., 2011, Du and Poo, 2004). Additionally, PLC-γ has been described to play a role in neurite outgrowth of sensory neurons (Sciarretta et al., 2010). Using an *in vitro* model of compartmentalized cultures of cortical neurons, we evaluated the change in the dendritic morphology induced by axonal stimulation of BDNF, and we investigated whether the presence of a well-known pharmacological aminosteroid PLC-γ antagonist, U-73122 (Bleasdale et al., 1990b, Smith et al., 1990, Smallridge et al., 1992), in the axons affects the retrograde signaling of BDNF using the protocol described in our previous work (Moya-Alvarado et al., 2023). In brief, we expressed EGFP in DIV 6 cortical neurons plated in microfluidic devices with 400 μm long microgrooves. We incubated a TrkB-Fc chimeric protein (Shelton et al., 1995) in the cell body compartment (CB) to neutralize the activity of endogenous BDNF released by neurons. To identify the neurons that projected their axons to the axonal compartment (AC), we used a fluorescent version of subunit B of cholera toxin (Ctb) (Escudero et al., 2019) that is internalized in axons and retrogradely transported up to the Golgi apparatus of neurons that have axons in the AC (Fig. 1A). To identify the somatodendritic domain of neurons, MAP2 immunofluorescence was performed. As we observed previously, the addition of BDNF to axons increased arborization (Fig. 1B and C), the primary dendrite numbers (Fig. 1D) and the branching point numbers (Fig. 1E) in rat cortical neurons (Moya-Alvarado et al., 2023). However, the presence of U-73122 in the AC prevented these effects of BDNF (Fig. 1 B-E), suggesting that PLC-γ is required for the long-distance axonal signaling of BDNF.

**Figure 1.**
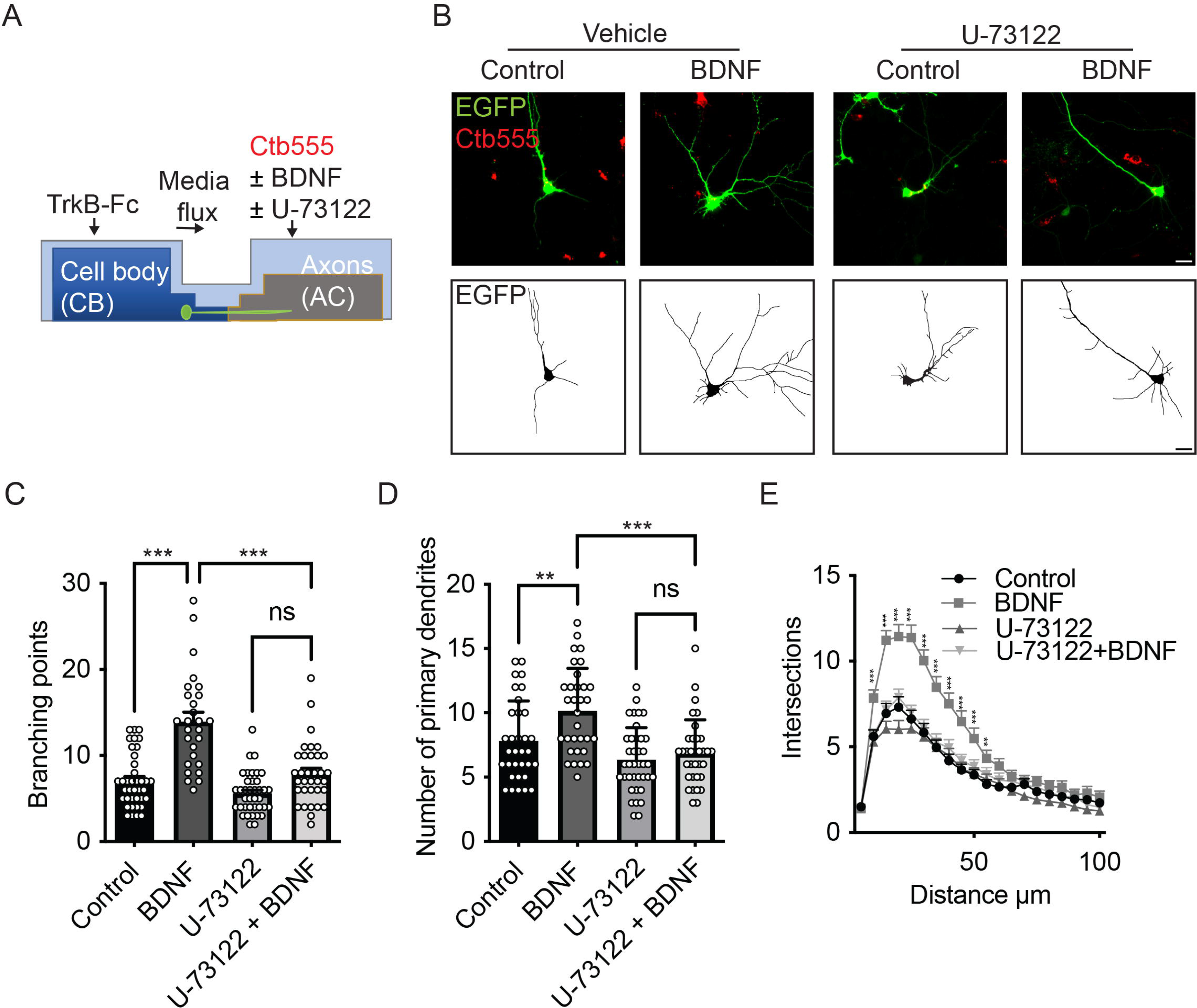
Axonal PLC-γ activity is required for dendritic arborization induced by BDNF. **(A)** Cortical neurons (DIV 6) were transfected with a plasmid expressing EGFP. The cell body compartment (CB) was incubated with TrkB-Fc (100 ng/mL). The axonal compartment (AC) was stimulated with BDNF (50 ng/mL) in addition to Ctb555 in the presence or absence of the PLC-γ inhibitor U73122 (5 µM). Treatments were performed for 48 hours. Finally, neurons were fixed, and immunofluorescence was performed against MAP2. **(B)** Representative images of the CB (red indicates Ctb555) of compartmentalized rat cortical neurons whose axons were treated with DMSO (control), U73122, BDNF or BDNF following preincubation with U73122. **(C-E)** Quantification of primary dendrites **(C)** and branching points **(D)** and Sholl analysis **(E)** for neurons labeled with EGFP/MAP2/Ctb555 for each treatment. Scale bar, 20 µm. n= 27-38 neurons from 3 independent experiments. The results are expressed as the means ± SEMs. **p< 0.01, ***p< 0.01. Statistical analysis was performed by one-way ANOVA followed by the Bonferroni correction for multiple comparisons (C-D). The Sholl’s analysis results were statistically analyzed by two-way ANOVA followed by the Bonferroni correction for multiple comparisons.

TrkB mutations in the PLC-γ docking region impair the phosphorylation of CREB and CaMKIV in non-compartmentalized cultures (Minichiello et al., 2002), suggesting that the activity of PLC-γ is required for CREB phosphorylation. Since we have previously described that axonal stimulation with BDNF promotes nuclear CREB phosphorylation (Moya-Alvarado et al., 2023, Bronfman et al., 2014) we tested whether the activity of PLC-γ was required in the cell body and/or axons for axonal BDNF-dependent CREB phosphorylation. To evaluate CREB phosphorylation, at DIV 5, we incubated Ctb555 in the AC overnight to identify all the neurons that projected their axons. On the next day, we added BDNF to the AC in the presence or absence of U-73122 in the AC or the CB. As we previously reported, axonal signaling of BDNF induced an increase in CREB phosphorylation (Moya-Alvarado et al., 2023). Interestingly, the presence of the PLC-γ inhibitor decreased CREB activation induced by axonal BDNF only when added to the AC (Fig. 2 A, B and C), having no effect when added to the CB (Fig. 2 C, D and E), suggesting that BDNF/TrkB axonal signaling to cell bodies requires PLC-γ activity mainly for propagation axonal signaling.

**Figure 2.**
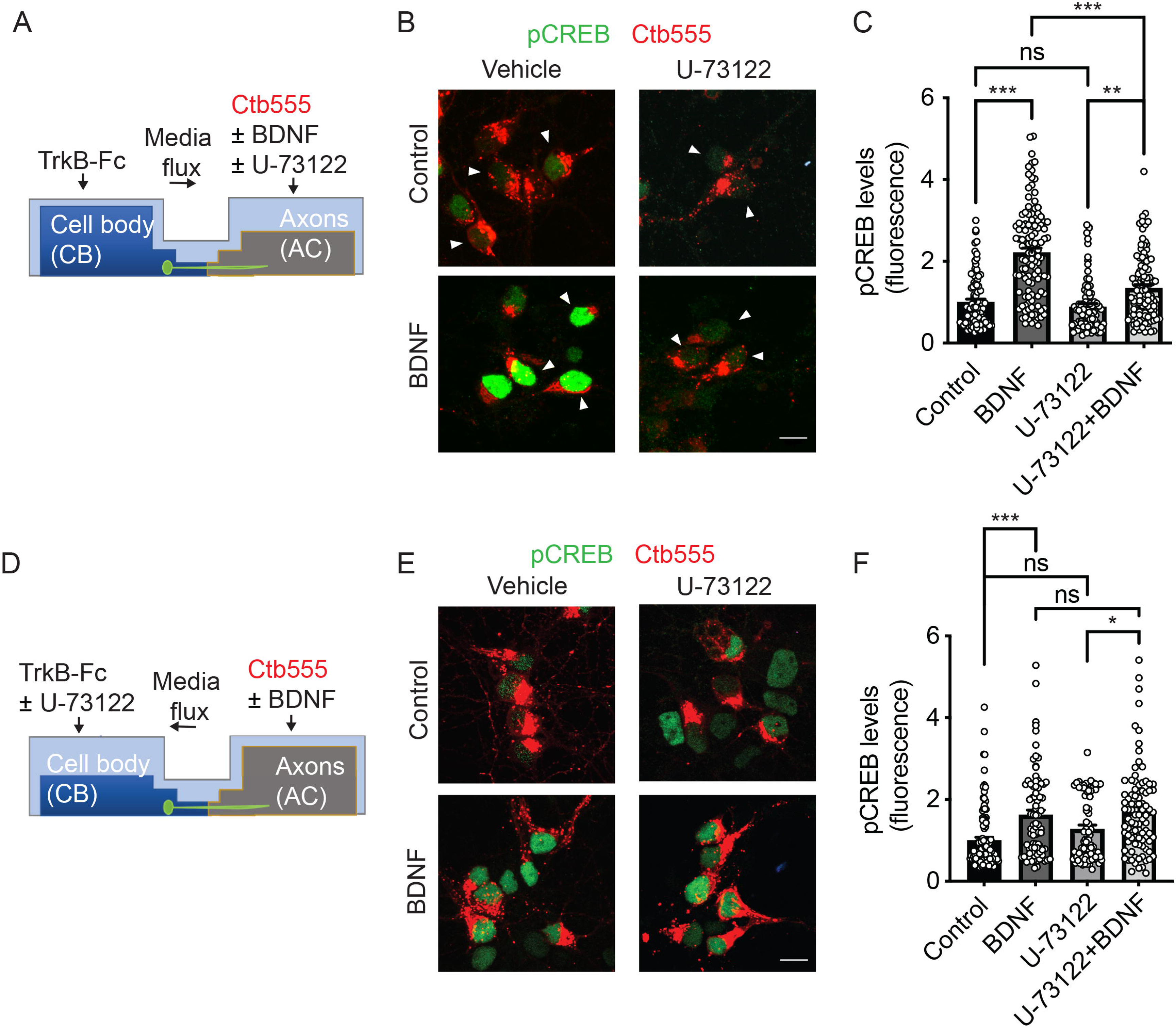
Axonal PLC-γ activity is required for somatodendritic CREB phosphorylation. **(A)** Schematic representation of the protocol used for stimulating neurons. DIV 5 cortical neurons were retrograde labeled with Ctb555 (in red) overnight. At DIV 6, the culture medium was changed to serum-free medium for 90 minutes in the presence or absence of U73122 (5 µM) in the AC and stimulated with BDNF (50 ng/mL) for 180 minutes in the AC in the presence or absence of U-73122 with the flux toward the AC. Finally, the cultures were fixed, and phosphorylated CREB (pCREB, S133) immunofluorescence was analyzed in cell bodies. **(B)** Representative figures of nuclear pCREB in neurons with or without BDNF stimulation labeled with Ctb555 (in red) added to axons in the presence or absence of axonal U-73122 as indicated in A’. Scale bar, 10 µm. **(C)** Quantification of pCREB in the nucleus of neurons labeled with Ctb555 (red) in each condition. n= 90-114 neurons from 3 independent experiments as shown in B. **(D)** Schematic representation of the protocol used for stimulating neurons. DIV 5 cortical neurons were retrograde labeled with Ctb555 (in red) overnight. At 6 DIV, the culture medium was changed to serum-free medium for 90 minutes in the presence or absence of U73122 (5 µM) in the CB with the flux toward the CB, and then the AC was incubated with BDNF (50 ng/mL) for 180 minutes. Finally, the cultures were fixed, and pCREB was analyzed in cell bodies **(E)** Representative figures of nuclear pCREB (in green) in neurons with or without BDNF stimulation labeled with Ctb555 (in red) added to axons in the presence or absence of U73122 in the CB. **(F)** Quantification of pCREB in the nucleus in neurons labeled with Ctb555 in each condition. n= 43-60 neurons from 2 independent experiments as indicated in D. The results are expressed as the means ± SEMs. **p< 0.01, ***p< 0.01. Statistical analysis was performed by one-way ANOVA followed by the Bonferroni correction for multiple comparisons.

### PLCγ is locally activated in the AC by BDNF and is required for the transport of BDNF-containing signaling endosomes

To test whether BDNF activates PLC-γ in axons, we treated the axons of cortical neurons with BDNF for 20 minutes and assessed the phosphorylation of PLC-γ by immunofluorescence staining in the microgrooves proximal to the AC. Immunofluorescence analyses with a phospho-specific antibody revealed that BDNF increases PLC-γ phosphorylation in the axons of cortical neurons, and immunostaining was prevented by treatment with K252a, an inhibitor of the tyrosine kinase activity of Trks (Tapley et al., 1992), confirming the specificity of the staining (Fig. 3A and B) (Tapley et al., 1992). The total protein level of PLC-γ was not changed by BDNF treatment (Fig. 3C and D and Fig.7C and D). The activation of PLC-γ by TrkB leads to an increase in intracellular Ca^2+^ (Zirrgiebel et al., 1995). To assess whether local stimulation of axons with BDNF increases cytosolic Ca^2+^ in a PLC-γ-dependent manner, we loaded the neurons with Fluo-4 AM, a cell permeant Ca^2+^ indicator (Cheng et al., 2017), and we recorded Fluo-4 AM fluorescence in the axons 1 minute after BDNF addition to study the immediate effect of BDNF on axonal Ca^2+^ levels. Incubation of axons with BDNF produced a single point Ca^2+^ release that generated an extended retrograde signal that completely covered the axons in the recording zone (Fig. 4A). Notably, not all the axons responded at the same time or with the same intensity. However, the presence of U-73122 completely abolished the increase in cytosolic Ca^2+^ induced by BDNF stimulation (Fig. 4A and B). We measured the speed of the retrograde Ca^2+^ signal considering the initial point of Ca^2+^ increase until the last point observable in the video recording; the measured speed was 4.47±0.15 μm/s (Fig. 4C), showing faster movement than that of dynein-dependent TrkB/BDNF retrograde trafficking (Xie et al., 2012). We previously showed that fluorescent BDNF undergoes retrograde transport with active TrkB receptors to the cell body in compartmentalized cultures of mouse cortical neurons (Moya-Alvarado et al., 2023). Consistent with the abovementioned results, the transport of BDNF-containing endosomes were dependent on both PLC-γ activity and the intracellular Ca^2+^ level (Fig. 5A, B, C and D).

**Figure 3.**
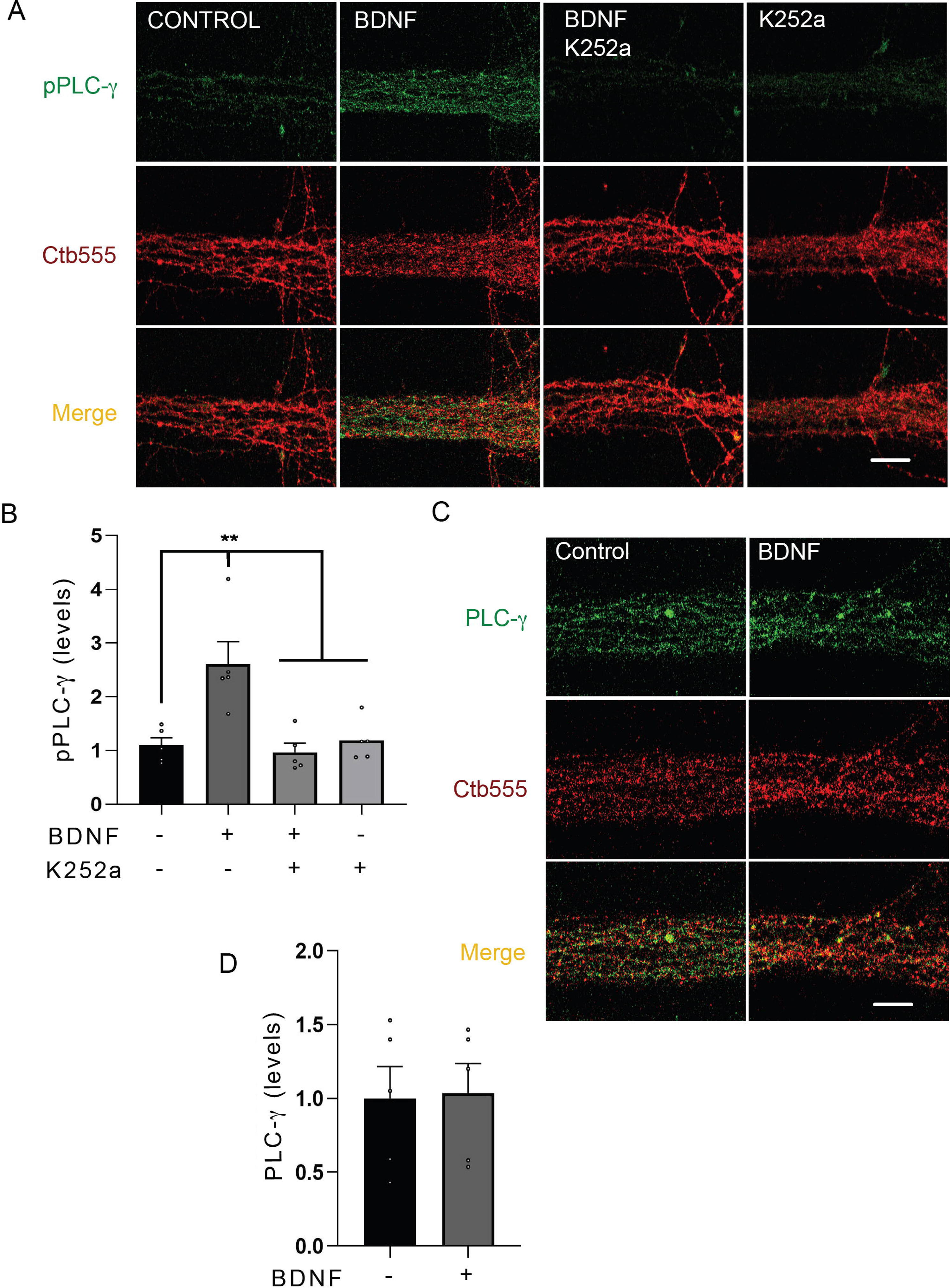
BDNF in axons promotes axonal PLC-γ phosphorylation. **(A)** Representative images of phosphorylated PLC-γ (pPLC-γ, pY783.28, green) in axons of compartmentalized rat cortical neurons left unstimulated (control), stimulated with 50 ng/mL BDNF for 20 minutes (BDNF) or stimulated with BDNF in the presence of 0.2 μM K252a (BDNF + K252a). Axons were labelled with Ctb555 overnight before treatment to assess correct compartmentalization of the culture. **(B)** Quantification of the immunofluorescence signal associated with pPLC-γ was performed in a rectangular ROI (as shown in the merged panel in A) drawn in the border between the beginning of the microgroove and the AC and continuing for 30 µm towards the microgroove that was delimited by Ctb555 fluorescence. n=5 unique chambers (the value in each chamber corresponds to the average of 5 different microgrooves) performed in five independent cultures. Statistical analysis was performed by one-way ANOVA followed by the Bonferroni correction for multiple comparisons. ** P<0.01. **(C)** Compartmentalized neurons were treated as described in A (but K252a was not used in this experiment). A representative image of total PLC-γ in the AC quantified as indicated in B used to assess pPLC-γ immunostaining is shown. **(D)** Quantification of the immunofluorescence signal associated with total PLC-γ. Statistical analysis was performed by Student’s t-test. No significant differences were found. Scale bar, 10 µm.

**Figure 4.**
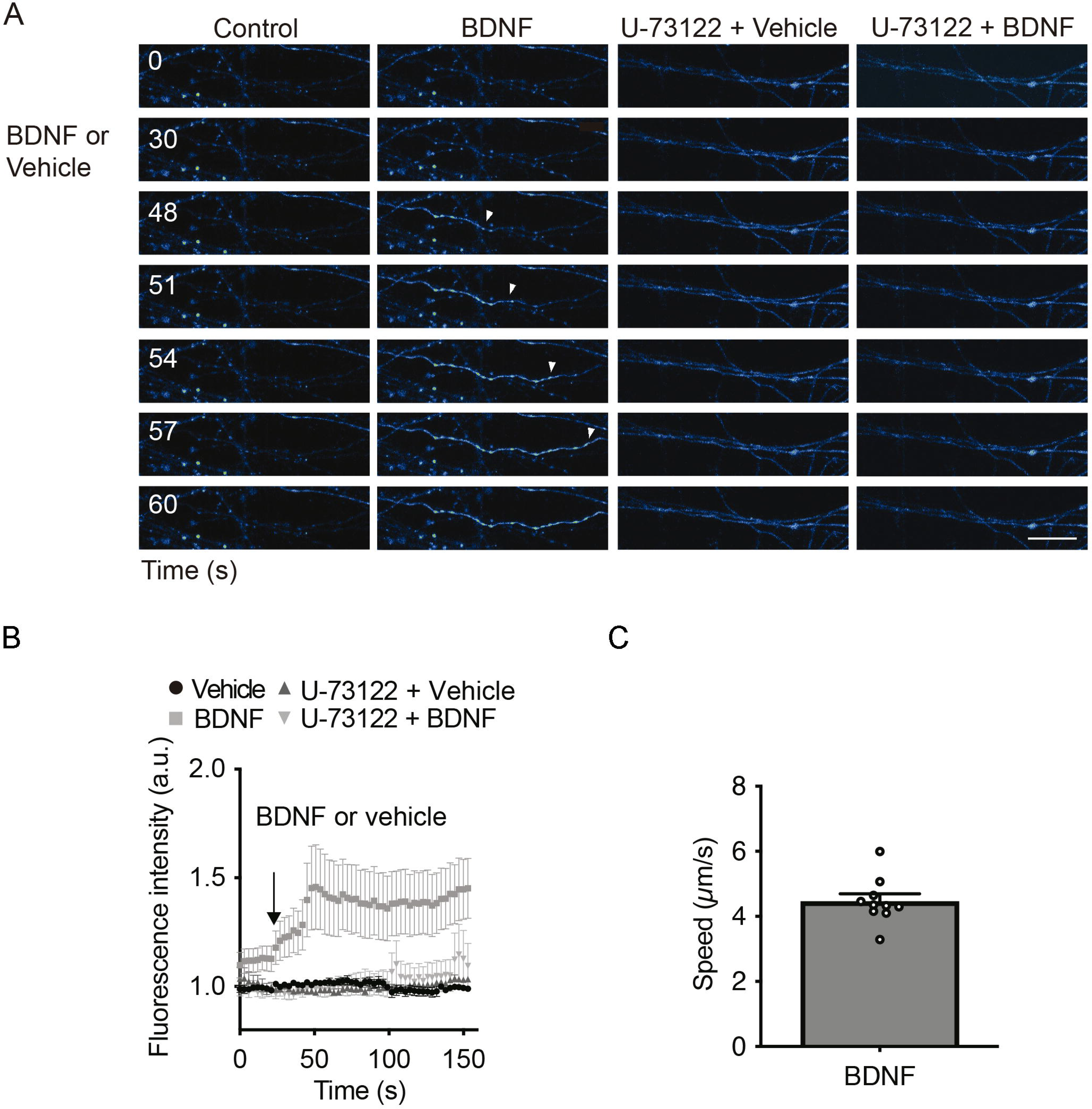
Axonal BDNF promotes an increase in intracellular Ca^2+^ in a PLC-γ-dependent manner. **(A)** Evaluation of Ca^2+^ signaling induced by BDNF. The change in fluorescence intensity associated with Fluo4-AM (2 μM) was used to measure the concentration change in cytosolic Ca^2+^. Representative images of compartmentalized cultures loaded with Fluo4-AM in the AC treated with vehicle or BDNF (50 ng/mL) in the AC with or without U73122 (5 µM) pretreatment. Live-cell imaging of each axonal field in the AC recorded before BDNF treatment (0 seconds) and during 60 seconds of BDNF treatment. Scale bar, 10 μm. **(B)** Mean Fluo4-AM fluorescence intensity (± SEM) for each treatment at different snapshot times. Normalization and statistical analysis for each treatment were performed using the corresponding mean ‘0 second’ basal fluorescence as a reference. n=8-12 axons from three independent compartmentalized cultures. Statistical analysis was performed by two-way ANOVA followed by the Bonferroni correction for multiple comparisons. **** P<0.0001. **(C)** Quantification of the velocity of Ca^2+^ back-propagation of the fluorescence signal associated with Fluo4-AM under BDNF conditions.

**Figure 5.**
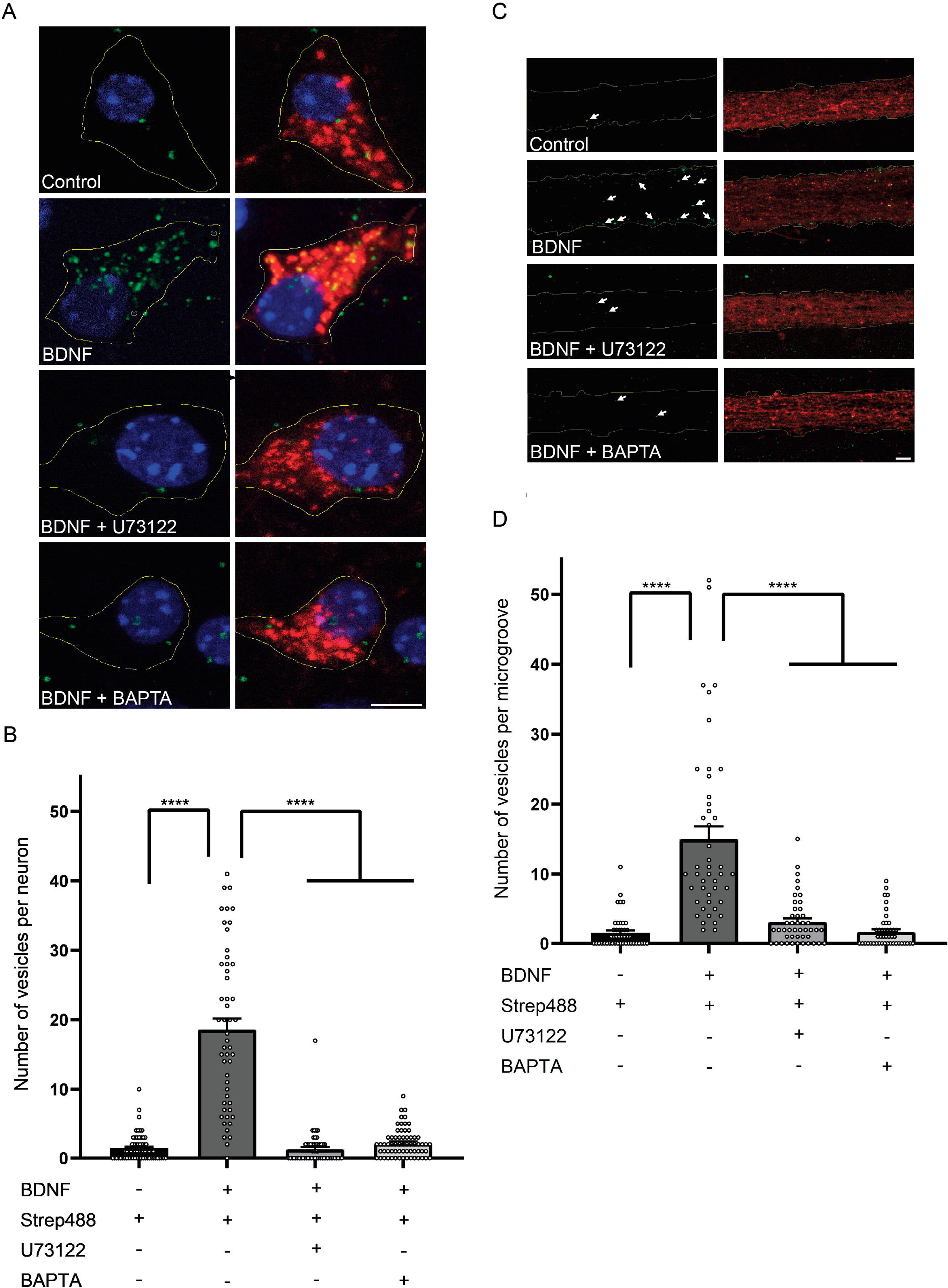
Axonal PLC-γ activity and intracellular Ca^2+^ are required for axonal BDNF-containing signaling endosome generation and retrograde transport to cell bodies. **(A)** DIV 7 compartmentalized mouse cortical neurons were retrograde labeled with Ctb555 (red) overnight. At DIV 8, the cell body compartment was treated with TrkB-Fc, and the AC was treated with DyLight 488-conjugated streptavidin alone (Control) or with biotinylated BDNF coupled to DyLight 488-conjugated streptavidin (f-BDNF, 150 ng/mL) for 6 hours in the absence (BDNF, green) or presence of 5 μM U-73122 (BDNF + U-173122) or 20 μM BAPTA, AM (BDNF + BAPTA). Then, the cells were fixed, mounted in Mowiol containing Hoechst (blue) and prepared for visualization. The circles in the BDNF panel indicate the smallest BDNF endosome considered for the analysis shown in B. Scale bar, 5 μm **(B)** Quantification of BDNF-positive vesicles in cell bodies of neurons containing Ctb555. Only vesicles larger than 20*10^-2^ μm^2^as shown in the BDNF treatment group by the circles in panel A were considered for the analysis. Forty-five neurons from three independent compartmentalized cultures were considered. Statistical analysis was performed by one-way ANOVA followed by the Bonferroni correction for multiple comparisons. **** P<0.0001. **(C)** Left panels, representative images of DyLight 488-conjugated streptavidin (green) associated fluorescence. White lines are selecting the region of the microgroove label with Ctb555 (red) shown in the right panels. White arrows indicate DyLight 488-conjugated streptavidin positive vesicles (control) or f-BDNF positive vesicles in the lower panels. Right panels, Representative images of axons (labeled with Ctb555, red) in microgrooves of neurons treated as described in A. Scale bar, 5 μm. **(D)** Quantification of BDNF-positive vesicles in microgrooves of axons containing Ctb555. Forty-five microgrooves from three independent compartmentalized cultures were considered for the analysis. Statistical analysis was performed by one-way ANOVA followed by the Bonferroni correction for multiple comparisons. **** P<0.0001.

### PLC-γ activity is required for BDNF-dependent TrkB internalization

Neurotrophin binding to Trks results in the internalization of the receptorLligand complex into endosomes, where the receptorLligand complex continues signaling (Beattie et al., 1996, Bronfman et al., 2014). PLC-γ regulates both epidermal growth factor receptor (EGFR) (Delos Santos et al., 2017) and TrkA internalization (Bodmer et al., 2011). Furthermore, TrkB internalization is regulated by neuronal activity and the influx of Ca^2+^, a second messenger downstream of PLC-γ activation (Du et al., 2003). In addition, the retrograde transport of BDNF-containing signaling endosomes depends on PLC-γ activity and intracelular Ca^2+^stores (Fig. 5), suggesting that PLC-γ activity regulates TrkB internalization and/or the retrograde transport of signaling endosomes. To specifically study whether PLC-γ activity is required for TrkB internalization, we performed an immunoendocytosis assay as described previously (Lazo et al., 2013) in non-compartmentalized primary cultures of cortical neurons. We transfected neurons with a plasmid expressing a TrkB receptor amino terminally tagged with a Flag epitope (Flag-TrkB) as previously described (Lazo et al., 2013). Forty-eight hours later, we treated the neurons with an anti-Flag antibody at 4°C. Next, we stimulated neurons with BDNF in the presence or absence of the PLC-γ inhibitor U-73122 (Fig. 6A). Although, there is a basal level of TrkB receptor internalization, BDNF increased TrkB internalization by approximately 50%, an effect that was significantly reduced when PLC-γ activity was inhibited (Fig. 6B and C). U-73122 did not influence the levels of Ctb555 internalization (Fig. 6D). These experiments were performed in rat cortical neurons, and we previously showed that BDNF has similar effects on mouse cortical neurons (Moya-Alvarado et al., 2023). Thus, we performed the same experiments shown in Fig.6 in mouse cortical neurons; however, we also included, in addition to the PLC-γ antagonist U-73122, the inactive analog U-73343 as a control (Smallridge et al., 1992, Smith et al., 1990, Bleasdale et al., 1990a). Similar to rat cortical neurons, mouse cortical neurons transfected with Flag-TrkB showed a basal level of recombinant TrkB internalization that was increased by 20 minutes of BDNF treatment (Fig. 7A and B). U-73122 almost completely abolished BDNF-dependent internalization of TrkB, as shown in rat cortical neurons (Fig. 6). However, in neurons treated with BDNF in the presence of U-73343, no reduction in receptor internalization was observed (Fig. 7A and B). Since U-73122 acts as an antagonist of PLC-γ downstream of receptor activation, these drugs should not influence BDNF-dependent phosphorylation of PLC-γ, as we observed in western blot analysis of mouse cortical neurons treated with or without BDNF in the presence or absence of U-73122 or U-73343 (Fig. 7C and D). Altogether, these results suggest that PLC-γ regulates BDNF-dependent TrkB internalization in cortical neurons.

**Figure 6.**
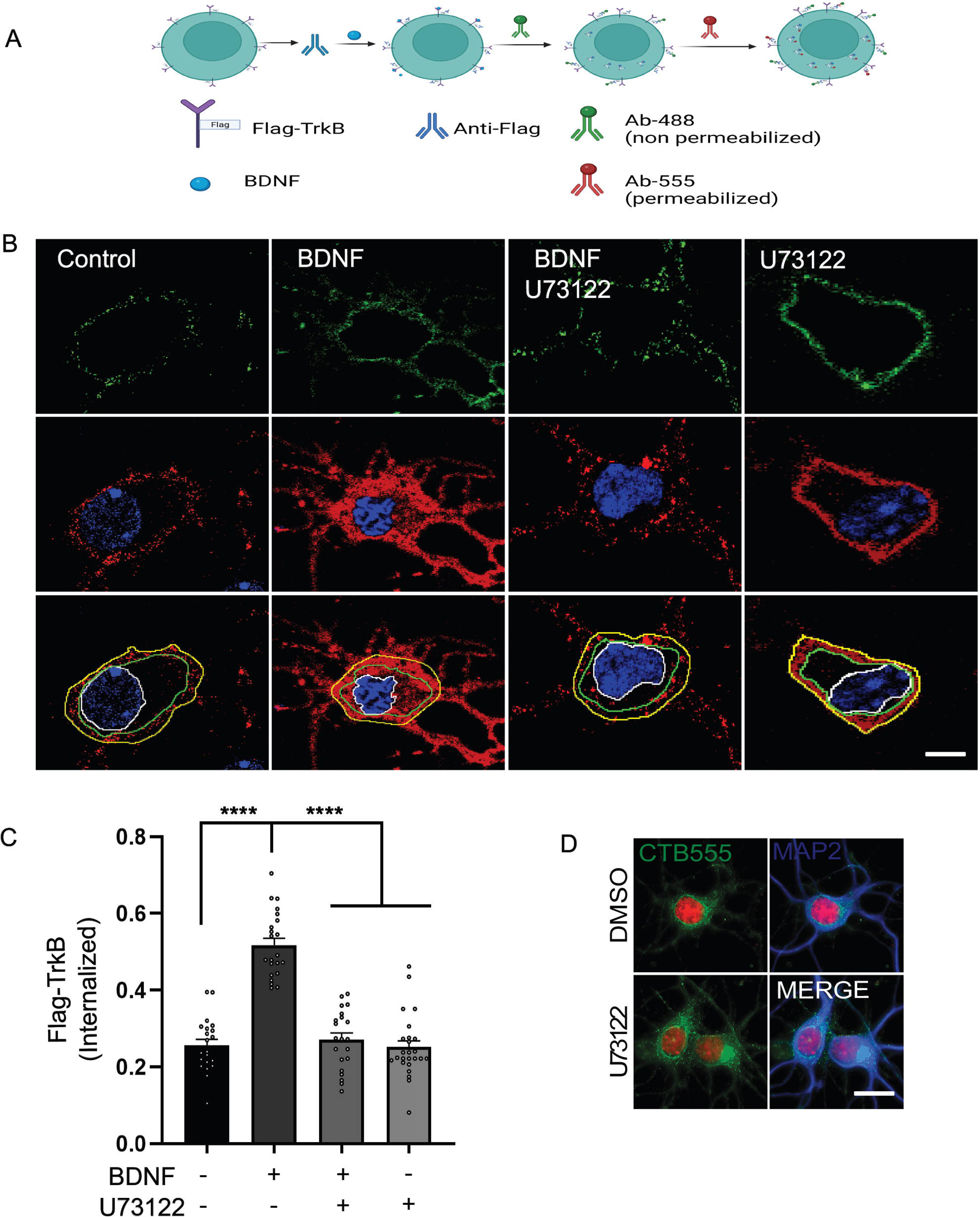
PLC-γ activity is required for TrkB internalization in rat cortical neurons. **(A)** Schematic representation of the immunoendocytosis protocol used to study Flag-TrkB internalization. Neurons (DIV 6) were transfected with a plasmid expressing Flag-TrkB. After 48 hours, the neurons were incubated with an anti-Flag antibody. Then, the neurons were treated with BDNF (50 ng/mL) in the presence or absence of U-73122 (5 µM) for 20 minutes at 37°C to induce endocytosis. Finally, the neurons were fixed, and the Flag epitope was detected by immunostaining. The plasma membrane-associated anti-Flag antibody was recognized with an anti-mouse Alexa 488 antibody (Ab-488, green) before cell permeabilization; then, cells were permeabilized, and the total Flag epitope was recognized with an anti-mouse Alexa 555 (red) antibody (Ab-555). **(B)** Representative images of the endocytosis of Flag-TrkB in control cells treated with BDNF, U-73122 or BDNF with U-73122. The upper figure shows plasma membrane-associated Flag-TrkB (in green), and the lower images show total Flag-TrkB in red; nuclei labeled with Hoechst are shown in blue. The yellow line delimits the whole cell and the total Flag-associated fluorescence (red), the space between the green line (delimiting membrane-associated Flag-TrkB) and the white line (delimiting the nucleus) is the cytosol-associated Flag-TrkB (in red). Scale bar, 5 µm. **(C)** Quantification of internalized Flag-TrkB was accomplished by dividing the fluorescence intensity associated with internalized Flag-TrkB (only the red fluorescence signal located between the green and white lines) by the fluorescence intensity associated with total intracellular Flag-TrkB under each treatment condition. **(D)** Representative image of Ctb555 endocytosis in the presence or absence of U-73122. Green, Ctb555; blue, Map2; red, Hoechst. Scale bar, 10 µm.

**Figure 7.**
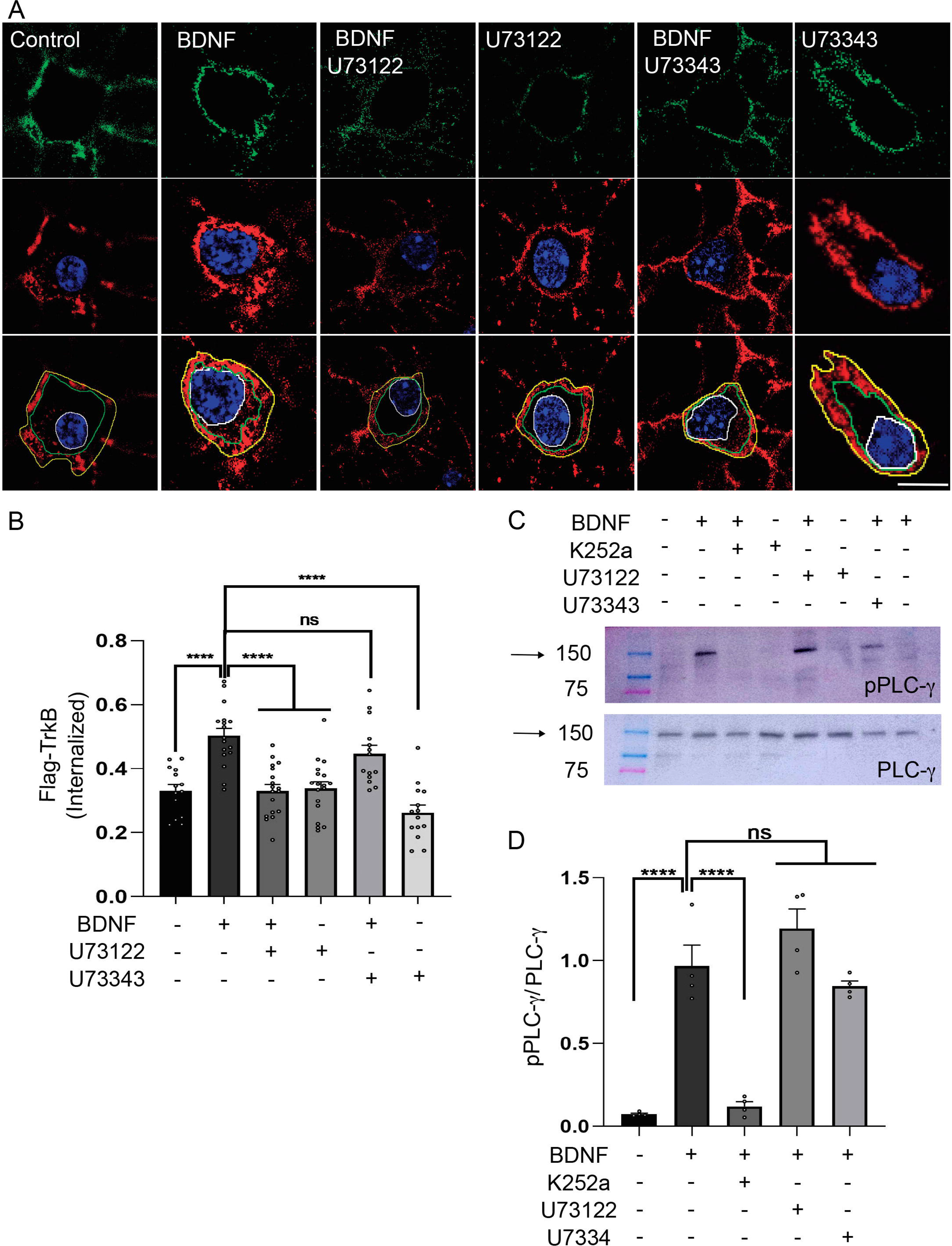
PLC-γ activity is required for TrkB internalization in mouse cortical neurons. Neurons (DIV 6) were transfected with a plasmid expressing Flag-TrkB. After 48 hours, the neurons were incubated with an anti-Flag antibody. Then, the neurons were treated with BDNF (50 ng/mL) in the presence or absence of U-73122 (5 µM) or with BDNF (50 ng/mL) in the presence or absence of U-73343 (5 µM) for 20 minutes at 37°C to induce endocytosis. Finally, the neurons were fixed, and the Flag epitope was detected by immunostaining. The plasma membrane-associated anti-Flag antibody was recognized with an anti-mouse Alexa 488 antibody (Ab-488) before cell permeabilization; cells were then permeabilized, and the total Flag epitope was recognized with an anti-mouse Alexa 555 antibody (Ab-555). **(A)** Representative images of Flag-TrkB endocytosis in control cells treated with BDNF, U-73122, U-73343, BDNF with U-73122 or BDNF with U-73343. The upper figure shows plasma membrane-associated Flag-TrkB (in green), and the lower images show total Flag-TrkB in red; nuclei labeled with Hoechst are shown in blue. The yellow line delimits the whole cell and the total Flag-associated fluorescence (red), the space between the green line (delimiting membrane-associated Flag-TrkB) and the white line (delimiting the nucleus) is the cytosol-associated Flag-TrkB (in red). Scale bar, 5 µm. **(B)** Quantification of internalized Flag-TrkB was accomplished by dividing the fluorescence intensity associated with internalized Flag-TrkB by the fluorescence intensity associated with total intracellular Flag-TrkB under each treatment condition (only the red fluorescence signal located between the green and white lines). **(C)** Western blot (WB) analysis of DIV 7 neurons treated with BDNF, K252a, U-73122, U73343, BDNF with K252a, BDNF with U-73122 or BDNF with U-73343. Upper panel, WB against phosphorylated PLC-γ (pPLC-γ). Lower panel, WB against total PLC-γ (PLC-γ). **D)** Quantification of pPLC-γ levels was accomplished by calculating the ratio of the signal associated with pPLC-γ to the signal associated with total PLC-γ (pPLC-γ/PLC-γ). Neurons were left untreated (control) or treated with BDNF (50 ng/mL) in the absence or presence of K252a, U-73122 or U-73343. n=4 independent experiments. The results are expressed as the means ± SEMs. ****p< 0.0001. Statistical analysis was performed by one-way ANOVA followed by the Bonferroni correction for multiple comparisons.

## DISCUSSION

We previously described that BDNF long-distance signaling from axons increases cellular responses leading to dendritic arborization (Moya-Alvarado et al., 2023). However, the downstream signaling pathways involved in this process are poorly understood. Here, we show that axonal PLC-γ activity is required for dendritic arborization and CREB phosphorylation induced by BDNF axonal signaling. Furthermore, the local axonal activation of PLC-γ regulates the intracellular Ca^2+^ increase, the transport of BDNF-containing signaling endosomes and TrkB endocytosis.

In this study, we have favored pharmacological inhibitors of PLC-γ/Ca^2+^ of downstream over genetic tools. These drugs used, allowed us to inhibit enzymatic or biological activities in a localized manner, that is, in the compartment of cell bodies or the axonal compartment of microfluidic cultures, since current genetic tools do not allow this type of evaluation. Undoubtedly, pharmacological inhibitors have an explicit limitation of selectivity. There is still room for the development of inhibitors that can be used to assess local activities in neuronal cells, such as optogenetic inhibition, as it has been used for RabGTPases and the TrkA receptor (Khamo et al., 2019, Nguyen et al., 2016, Cornejo et al., 2020)

We have previously described that somatodendritic but not axonal signaling of PI3K is required for cellular responses associated with BDNF axonal signaling (Moya-Alvarado et al., 2023). Here, we showed that the axonal activity of PLC-γ is required for dendritic arborization induced by BDNF in axons. Interestingly, an *in vivo* experiment with RGCs of the *Xenopus laevis* optic tectum showed that tectal stimulation with BDNF promotes a potentiation of retinotectal synapses in a retrograde-dependent manner. This process was dependent on the activity of TrkB and PLC-γ, suggesting that there are synaptic modifications that may be causally linked to structural modification of RGC dendrites after hours of BDNF stimulation in the axons (Du and Poo, 2004, Lom et al., 2002b). Consistently, we showed that BDNF addition to the axons of cortical neurons increased dendritic branching (Moya-Alvarado et al., 2023), although synaptic contacts did not develop due to the limited time in culture (6-8 days). In future experiments, it will be important to evaluate whether BDNF axonal signaling promotes synaptic strengthening in mature neurons using compartmentalized cultures.

In the same context, we observed that axonal but not cell body activity of PLC-γ is required for CREB phosphorylation induced by BDNF axonal signaling, suggesting that PLC-γ has an axonal local role in cortical axons. It is likely that the initial steps of signaling endosome generation are regulated. Inhibition of PLCγ in cell bodies did not reduce CREB activation induced by the addition of axonal BDNF, indicating that PLC-γ is required for the initiation and propagation of axonal signaling endosomes but does not participate in the CREB phosphorylation induced by BDNF axonal signaling after this signaling has reached the cell body. It has been shown that mutation of the TrkB docking site for PLC-γ downregulates BDNF-induced activation of CREB in non-compartmentalized neuronal cultures (Minichiello et al., 2002). Therefore, it is possible that BDNF signaling from different locations globally contributes to CREB activation.

In non-compartmentalized cultures, bath application of BDNF promotes a strong increase in the frequency of global Ca^2+^ transients (Lang et al., 2007, He et al., 2005), but local BDNF application induces a point-spread signal in dendrites (Lang et al., 2007). We observed that BDNF axonal stimulation promoted an increase in intracellular Ca^2+^, and like the observation reported in dendrites, we observed a single Ca^2+^ point-spread that generated an extended retrograde signal that covered the axons along the recording zone. Since TrkB is on the surface of the axons, it is intriguing that Ca^2+^ waves are generated in apparently random locations along the axons. Of note, we observed that the velocity of the Ca^2+^ waves was faster than the average velocity of BDNF-QD (1.11±0.05 µm/s) (Xie et al., 2012) and (1.5 ±0.3 µm/s) for GFP-TrkB (Goto-Silva et al., 2019), suggesting that this Ca^2+^ increase is independent of the retrograde transport of the signaling endosome. Interestingly, it was recently shown that the Ca^2+^-dependent protein phosphatase calcineurin is in the cytosolic part of signaling endosomes, where it dephosphorylates huntingtin in a Ca^2+^-dependent manner to favor the retrograde transport of TrkB endosomes. This suggests that the Ca^2+^ waves induced by BDNF can promote the retrograde trafficking of axonal TrkB endosomes (Scaramuzzino et al., 2022).

In sensory neurons, back-propagation of Ca^2+^ at an injury site has been reported along the axon toward the cell soma (Cho et al., 2013). This process regulates dual leucine zipper kinase (DLK), which activates c-Jun NH2 terminal kinase (JNK3); JNK3 is linked to axonal transport via JNK-interacting protein JIP3, a protein that interacts with kinesin and dynactin (Cavalli et al., 2005, Rishal and Fainzilber, 2014). Interestingly, it has been shown that JIP3 is an adaptor for the anterograde transport of TrkB mediated by kinesin 1 (Huang et al., 2011). However, the increase in intracellular Ca^2+^ in axons increases the affinity of JIP3 for dynactin compared to kinesin (Cavalli et al., 2005, Rishal and Fainzilber, 2014). Therefore, it is possible that the increase in intracellular Ca^2+^ ions mediated by BDNF/TrkB in axons increases the TrkB/JIP3/dynactin/dynein complex to promote the retrograde transport of the TrkB receptor.

The propagation of Ca^2+^ waves may not only facilitate the retrograde trafficking of endosomes but also increase the probability of TrkB internalization along the axon. We found that PLC-γ activity was required for TrkB internalization. PLC-γ has been described as a regulator of TrkA and EGFR endocytosis, and two different mechanisms have been reported for this PLC-γ-dependent internalization. For TrkA, Ca^2+^ contributes to enhanced clathrin-mediated endocytosis in neurons due to calcineurin-dependent dephosphorylation of dynamin 1 (Bodmer et al., 2011). On the other hand, calcineurin and dynamin 1 are dispensable for the internalization of EGFR. In this case, Ca^2+^ ions recruit synaptojanin 1 (Sjn1) to clathrin-coated pits in a PKC-dependent manner induced by the activation of PLC-γ (Delos Santos et al., 2017).

In summary, this study reveals a specific function for PLC-γ signaling in the AC of cortical axons, regulating the signaling of BDNF/TrkB from distal axons to the cell body to increase CREB-dependent dendritic branching. Several processes remain to be clarified in future studies, including the mechanism accounting for the PLC-γ/Ca^2+^-dependent regulation of TrkB in axons and the contribution of TrkB/Ca^2+^ to the trafficking of signaling endosomes.

## METHODOLOGY

### Primary culture of cortical neurons

Embryonic cortical neurons were obtained from mice (C57Bl/6J) and rats (*Rattus norvegicus*) (embryonic days 16–18) housed in the animal facilities of our institutions. Pregnant animals were euthanized under deep anesthesia according to bioethical protocols approved by the Bioethics Committee of the Pontificia Universidad Catolica (protocol ID:180822013) de Chile and Universidad Andres Bello (act of approval 022/2019 and 009/2022).

Rat and mouse cortical tissues were harvested and dissociated into single cells in Hank’s balanced salt solution (HBSS; Thermo-Fisher, cat# 14025134). After disaggregation, the neurons were resuspended in modified Eagle’s medium (MEM) supplemented with 10% horse serum (HS) (MEM/HS, Thermo-Fisher, cat# 16050122) and seeded in microfluidic chambers at a low density (40-50 × 10^3^ cells/chamber) or in mass culture at a density of 35 × 10^3^ cells/well on 12 mm coverslips or 1.5 × 10^6^ cells/60 mm plate. After 4 h, the culture medium was replaced with neurobasal medium (Thermo-Fisher, cat# 21103049) supplemented with 2% B27 (Life Technologies, cat# 17504044), 1x GlutaMAX (Thermo-Fisher, cat# 35-050-061) and 1x penicillin/streptomycin (Thermo-Fisher, cat# 15140-122). The proliferation of nonneuronal cells was limited using cytosine arabinoside (0.25 µg/mL AraC; SigmaLAldrich, cat# C1768) when MEM/HS was replaced with neurobasal medium (Shimada et al., 1998, Taylor et al., 2003, Moya-Alvarado et al., 2023).

### Microfluidic devices

We thank the lab of Professor Eran Perlson at the Department of Physiology and Pharmacology of the Faculty of Medicine of Tel Aviv University for providing the molds to prepare the compartmentalized chambers (Gluska et al, 2016). The microfluidic chambers were prepared with SYLDGARD^TM^ 184 silicone elastomer base (Poirot, cat# 4019862) according to the manufacturer’s instructions. Two days before primary cultures, glass coverslips (25 mm) were incubated with poly-D-lysine (0.1 mg/mL, Corning, cat# 354210). The next day, poly-D-lysine was washed, and a microfluidic chamber with a 400 µm microgroove was placed on the coverslip. Then, laminin (2 µg/mL in water, Invitrogen, cat# 23017015) was added to the chamber. The same day, the primary culture laminin solution was changed to DMEM/HS medium (Dulbecco minimum essential medium supplemented with 10% horse serum, 1x GlutaMAX and 1x antibiotic/antimycotic, Thermo-Fisher, cat# 15240062).

### Quantification of BDNF-induced dendritic arborization in rat cortical neurons

Cortical neurons (DIV 6) were transfected with 0.5 µg of the plasmid containing EGFP (CA, USA) using 0.8 µL of Lipofectamine 2000 (Invitrogen, cat# 11668-019) in 30 µL of Opti-MEM (Thermo-Fisher, cat# 11058021). After 2 hours, the Opti-MEM medium was replaced with neurobasal medium, and the cells were incubated for 1 hour. The CB was incubated with neurobasal medium supplemented with TrkB-Fc (100 ng/mL, B&D system, 688TK) for all treatments. The drugs were incubated in the CB or AC. In the AC, U-73122 (5 µM, SigmaLAldrich cat# U6756) was used. After 1 hour, BDNF (50 ng/mL, Alomone, cat# B-250) was added to the AC together with fluorescent subunit B of cholera toxin (Ctb, 1 µg/mL, Thermo-Fisher, cat# C34777). After 48 hours, neurons were washed with PBS 37°C and then fixed with fresh 4% PFA-PBS at 37°C for 15 minutes (paraformaldehyde (SigmaLAldrich, cat# 158127) in PBS). Then, the chamber was removed, and the neurons were permeabilized and blocked with 5% BSA (Jackson, cat# 001-000-162) and 0.5% Triton X-100 (SigmaLAldrich, cat# 234729) in PBS and then incubated with anti-MAP2 (1:500, Merck-Millipore, cat# AB5622) in incubation solution (3% BSA with 0.1% Triton X-100 in PBS). After washes were performed (3x buffer), the neurons were treated with a donkey anti-mouse Alexa 647 antibody (1:500, Molecular Probes, cat# A-31571) in incubation solution and mounted for fluorescence microscopy visualization using Mowiol 4-88 (Calbiochem, cat# 475904).

Dendritic arborization was analyzed in cortical neurons labeled with Ctb and labeled for EGFP and MAP2. Primary dendrites, branching points and Sholl’s analysis (Sholl, 1953) were quantified (see below). For visualization, confocal microscopy using a Nikon Eclipse C2 confocal microscope equipped with a digital camera connected to a computer with Software NIS-Elements C was used. Five to seven optical sections (0.5 µm thick) from the whole cells, using the EGFP associated fluorescence, were acquired using a 60x objective at a resolution of 1024×1024 pixels along the z-axis. The z-stacks were integrated, and the images were segmented to obtain binary images. Ten concentric circles with increasing diameters (5 µm each step) were traced around the cell body, and the number of intersections between dendrites and circles was counted and plotted for each diameter. Analysis was performed using the ImageJ plugin developed by the Anirvan Gosh Laboratory (http://biology.ucsd.edu/labs/ghosh/software). The total primary dendrites and branching points of all dendrites were manually counted from the segmented images.

### Evaluation of CREB protein phosphorylation by immunofluorescence staining in rat cortical neurons

To evaluate the levels of phosphorylated CREB (pCREB) in the nucleus, experiments were performed as indicated in (Moya-Alvarado et al., 2023). In brief, rat cortical neurons (DIV 6) were incubated with Ctb555 (Ctb, 1 µg/mL) overnight. Next, cortical neurons (DIV 7) were incubated with neurobasal medium containing TrkB-Fc (100 ng/mL) in the cell body compartment. To inhibit PLC-γ, we used the drug U-73122 (5 µM) in the AC or cell body compartment depending on the experimental design. After 1 hour of inhibitor pretreatment, BDNF (50 ng/mL) was added to the AC. After 180 minutes, samples were fixed with 4% PFA supplemented with a phosphatase inhibitor (1x) for 15 minutes. Permeabilization and blocking were performed with bovine serum albumin (BSA, 5%), Triton X-100 (0.5%) for 1 hour in PBS. The samples were incubated with antibodies overnight at 4°C in the presence of BSA (3%) and Triton X-100 (0.1%) in PBS. An anti-phospho-CREB antibody (1:500, Cell Signaling cat# 9198) was used. The samples were incubated with secondary antibodies for 90 minutes in PBS containing BSA (3%) and Triton X-100 (0.1%). Finally, the samples were incubated with Hoechst 33342 (5 µg/mL, Invitrogen, cat# 62249) and mounted in Mowiol 4-88. Neurons were visualized by confocal microscopy using a Nikon Eclipse C2 confocal microscope equipped with a digital camera connected to a computer with NIS-Elements C software. Five to seven optical sections (0.5 µm thick) from the whole cells of Ctb555-positive neurons were acquired using a 60x objective at 1024×1024 pixel resolution along the z-axis.

### Evaluation of BDNF-induced phosphorylation of PLC-γ in axons of rat cortical neurons

To measure the levels of phosphorylated PLC-γ (pPLC-γ) in axons after BDNF treatment, rat cortical neurons (DIV 6) in compartmentalized cultures were incubated with Ctb555 (1 µg/mL) overnight. The next day (DIV 7), the neurons were incubated with neurobasal medium containing TrkB-Fc (100 ng/mL) in the cell body compartment, and the AC was pre-incubated for 1 hour with vehicle or 0.2 µM K252a (Tocris cat# 1683) (Tapley et al., 1992). Then, BDNF (50 ng/mL) was added to the AC in the presence or absence of K252a. After 20 minutes, the samples were fixed with 4% PFA/4% sucrose supplemented with a phosphatase inhibitor (Pierce, cat# A32957) for 15 minutes at room temperature (RT). Permeabilization and blocking were performed with PBS containing BSA (5%) and Triton X-100 (0.5%) for 1 hour. The samples were incubated with antibodies overnight at 4°C in PBS containing BSA (3%) and Triton X-100 (0.1%). Anti-phospho-PLC-γ (pY783.28, 1:200, Santa Cruz cat# sc-136186) and anti-PLC-γ1 (PLC-γ1, 1:200, Santa Cruz, cat# sc-7290) antibodies were used. The samples were incubated with secondary antibodies for 90 minutes in PBS containing BSA (3%) and Triton X-100 (0.1%). Finally, the samples were mounted in Mowiol 4-88 supplemented with Hoechst 33342 (5 µg/mL, Invitrogen, cat# 62249). To measure the level of total PLC-γ, K252a was not used.

Axons were visualized by spinning disk microscopy (Olympus DSU Spinning Disc) on a system equipped with a motorized stage and a digital camera connected to a computer running Olympus cellSens dimension software V 1.14. Whole axons positive for Ctb555 were visualized using an oil immersion (60x) objective at a resolution of 1024×1024 pixels along the z-axis. For quantification, the region of interest (ROI) was defined as the microgroove region nearest the AC. Five images were acquired per chamber (5 microgrooves were evaluated per chamber). For quantification, Image J 1.53t software was used. From each sample, the image with the highest Ctb55 signal along the z-axis was selected. The image was separated in two channels (red, Ctb555 and green, pPLC-γ or total PLC-γ) and converted to 8 bits for analysis. Then, the ROI was defined as a rectangle 15 µm high and 30 µm long, and the mean fluorescence intensity was calculated (total fluorescence intensity divided by 450).

### Measurement of axonal intracellular Ca^2+^

The changes in intracellular Ca^2+^ concentrations were examined by Fluo4-AM staining. Neurons were incubated with Fluo4-AM (2 µM, Invitrogen cat# F14201) in neurobasal medium for 30 minutes at 37°C and then incubated for another 15 minutes after rinsing with HBSS. Ca^2+^ imaging was performed by confocal microscopy using a Nikon Eclipse C2 confocal microscope equipped with a digital camera connected to a computer with NIS-Elements C software. The cells were excited with a laser at 488 nm, and the intensity of the fluorescence was collected at 525 nm as the Fluo4-AM signal. The images were collected before and after BDNF treatment (30 seconds after initial video recording), and the fluorescence intensity was analyzed with ImageJ. To measure the Ca^2+^ in the axonal terminals in the presence of U73122 (5 µM), neurons were preincubated in the axonal terminal with U73122 after Fluo4-AM loading.

### Quantification of BDNF-containing signaling endosomes in compartmentalized cultures of mouse cortical neurons

To prepare fluorescent BDNF (f-BDNF), commercially available biotinylated BDNF (Alomone Labs, cat# B-250-B) was coupled to DyLight 488-conjugated streptavidin (Invitrogen, cat# 21832) by incubation at 37°C for 20 minutes; the reaction consisted of 6 µL of a 6 µM solution of DyLight 488-conjugated streptavidin and 2 µL of a 6 µM solution of biotinylated BDNF diluted in 72 µL of neurobasal medium supplemented with 0.1% BSA (molar range 1:1). Then, this solution was diluted in the same buffer to a final concentration of 130 ng/mL BDNF as described in (Moya-Alvarado et al., 2023).

Experiments with compartmentalized mouse cortical neurons were performed as described in (Moya-Alvarado et al., 2023). In brief, mouse cortical neurons (DIV 7) were incubated with Ctb555 (1 µg/mL) overnight. Next, neurons (DIV 8) were incubated with neurobasal medium containing TrkB-Fc (100 ng/mL) in the CB compartment and with vehicle (DMSO), U-73122 (5 µM) or 20 µM BAPTA, AM (Invitrogen, cat# B6769) for 1 hour at 37°C in the AC. Then, f-BDNF was added to the AC for 6 hours in the presence or absence of the inhibitors. Then, the drugs were removed, and the neurons were fixed in 4% PFA/4% sucrose in PBS for 18 minutes, washed in PBS and distilled water and mounted in Mowiol 4-88 supplemented with Hoechst 33342 (5 µg/mL). Cell bodies and axons in the microgrooves were visualized using a confocal microscope (Leica SP8). Fifteen to twenty optical sections (0.5 µm thick) from the whole cell body or axons positive for Ctb555 were acquired using an oil immersion (63x) objective at a resolution of 1024×1024 pixels along the z-axis. For cell bodies and axons, a 5x or 2.5x digital zoom setting was used, respectively. Fifteen images, containing 1-4 cells or at least one microgroove, were acquired per condition in each experiment. For quantification, Image J 1.53t was used. To create one image, the maximum projection in each channel was generated using the z-stack images, and the background was reduced in the green channel (f-BDNF). The same procedure was performed for all conditions. All f-BDNF (green)-positive vesicles in each cell were selected, and the diameters were calculated. Only vesicles with an area greater than 20*10^-2^ µm^2^ were used for quantification, and the results are expressed as the number of BDNF-positive vesicles per cell or microgroove.

### Flag-TrkB immunoendocytosis assay to evaluate the effect of PLC-γ activity on the internalization of TrkB

Rat or mouse cortical neurons (DIV 6) were transfected with 0.5 µg of the Flag-TrkB plasmid (a gift from Prof. Francis Lee, Weill Cornel Medicine, New York, NY, US) using 1 µL of Lipofectamine 2000 in 100 µL of Opti-MEM per cover. After 48 hours, cortical neurons (DIV 8) were pretreated for 1 hour with TrkB-Fc (100 ng/mL) at 37°C and were then incubated on ice for 10 minutes and treated with an anti-Flag antibody (1:500, SigmaLAldrich, cat# F3040) for 45 minutes on ice in the presence or absence of 5 µM U-73122 or 5 µM U-74333 (U-7433 was used only in the experiments with mouse cortical neurons). Then, the cortical neurons were washed briefly with warm (RT) neurobasal medium and incubated with BDNF (50 ng/mL) in the presence or absence of U-73122 for 20 minutes. The neurons were washed twice with RT PBS and fixed in 4% PFA/4% sucrose in PBS for 15 minutes at RT. Then, the cells were washed twice with PBS and incubated with a donkey anti-mouse Alexa 488 antibody (1:300, Life Technologies, cat# A-2120) without permeabilization for 1 hour. Then, the samples were permeabilized and blocked with a solution of PBS supplemented with 5% BSA and 0.5% Triton X-100 for 1 hour at room temperature. After 1 PBS wash, the samples were incubated with a donkey anti-mouse Alexa 555 antibody (1:300, Invitrogen, cat# A-31570) diluted in PBS supplemented with 1% BSA and 0.1% Triton X-100 overnight at 4°C. Finally, the samples were washed in PBS and distilled water and mounted in Mowiol 4-88 supplemented with Hoechst 33342 (5 µg/mL).

Neurons were visualized using a confocal microscope (Leica SP8). Twelve to fifteen optical sections (0.5 µm thick) from whole neurons were acquired using an oil immersion (63x) objective at a resolution of 1024×1024 pixels along the z-axis. For quantification, the Image J 1.53t software was used. To quantify the internalization of Flag-TrkB receptors, the optical section passing through the middle of the soma having the most intense red signal was evaluated (anti-mouse Alexa 555). The image was separated in two channels (red, anti-mouse Alexa 555 and green, anti-mouse Alexa 488) and converted to 8 bits for analysis. To limit the red signal associated with the plasma membrane, the green signal corresponding to the anti-mouse Alexa 488 antibody (Flag epitope present in the plasma membrane) was used. Internalized Flag-TrkB was indicated by the red signal associated with the anti-mouse Alexa 555 antibody within the region of the cell not including areas associated with the plasma membrane. The total Flag-TrkB content was represented by the total signal associated with the anti-mouse Alexa 555 antibody. To normalize the level of Flag-TrkB expression, the intensity of anti-mouse Alexa 555 fluorescence associated with internalized TrkB was divided by the total fluorescence intensity associated with the anti-mouse Alexa 555 antibody (see Fig. 6 and 7).

### Western blot analysis of phosphorylated and total PLC-γ

One million five hundred mouse cortical neurons were seeded in 60 mm plates. After DIV 8, the neurons were treated for one hour with TrkB-Fc (100 ng/mL) in the presence or absence of the inhibitor U-73122 (5 µM) or U-74333 (5 µM). Then, the plates were treated with BDNF (50 ng/mL) in the presence or absence of the inhibitors. After treatment, the cells were placed on ice and washed in cold PBS twice, and lysates were prepared with RIPA buffer (150 mM sodium chloride, 1.0% Triton X-100, 0.5% sodium deoxycholate, 0.1% sodium dodecyl sulfate, 50 mM Tris (pH 8.0)). The homogenates were placed in an Eppendorf tube and centrifuged for 20 minutes at 12000 rpm and 4°C in a benchtop centrifuge. Then, the supernatant was placed in an Eppendorf tube on ice, and protein concentrations were quantified. To detect pPLC-γ and total PLC-γ, 20 µg of protein from each sample was loaded into an SDS-PAGE gel and transferred onto PVDF membranes. Membranes were activated with methanol and blocked with 5% BSA (Jackson ImmunoResearch, cat# 001-000-162) |in TBST (150 mM NaCl, 20 mM Tris-HCl (pH 7.4), 0.1% Tween® 20) for 1 hour. Incubation with the mouse anti-pPLC-γ (1:1000) primary antibody was performed in TBST overnight at 4°C. After three washes with TBST, the membranes were treated with a 1:10000 dilution of a mouse anti-HRP antibody (BioRad, cat# 1706516) for 2 hours and developed with a bioluminescence kit (Biological Industries, cat# 20-500-1000B) using an imaging system (GE Healthcare Life Science, ImageQuant™ LAS 500). Then, the membranes were stripped (Restore™ PLUS Western Blot Stripping Buffer, Thermo Scientific, cat# 4630) for 15 minutes at RT and blocked with 5% nonfat milk (Rockland, cat# b501-0500) in TBST for 1 hour. Then, the membrane was incubated with a mouse anti-PLC-γ antibody (1:500, Santa Cruz, cat# sc-7290) overnight. After three washes with TBST, the membrane was treated with a 1:10000 dilution of the mouse-anti-HRP antibody and developed as indicated above. To quantify the bioluminescence associated with each band, the “Gels” plugin in ImageJ 1.53t was used.

### Statistical analysis

The results are expressed as the averages ± standard errors of the mean (SEMs). Sholl’s analysis curves were compared with two-way repeated-measures ANOVA followed by the Bonferroni correction for multiple comparisons. Student’s t test or one-way ANOVA followed by the appropriate multiple comparisons test was performed depending on the number of groups used in each experiment. The details about the specific test used, level of significance and number of replicates are indicated in each figure legend. Statistical analyses were performed using GraphPad Prism 7 (Scientific Software).

## REFERENCES

Alonso, M., Vianna, M. R., Depino, A. M., Mello E Souza, T., Pereira, P., Szapiro, G., Viola, H., Pitossi, F., Izquierdo, I. & Medina, J. H. 2002a. BDNF-triggered events in the rat hippocampus are required for both short- and long-term memory formation. Hippocampus, 12, 551–60.

Alonso, M., Vianna, M. R., Izquierdo, I. & Medina, J. H. 2002b. Signaling mechanisms mediating BDNF modulation of memory formation in vivo in the hippocampus. Cell Mol Neurobiol, 22, 663–74.

Arimura, N., Kimura, T., Nakamuta, S., Taya, S., Funahashi, Y., Hattori, A., Shimada, A., Menager, C., Kawabata, S., Fujii, K., Iwamatsu, A., Segal, R. A., Fukuda, M. & Kaibuchi, K. 2009. Anterograde Transport of TrkB in Axons Is Mediated by Direct Interaction with Slp1 and Rab27. Developmental Cell, 16, 675–686.

Beattie, E. C., Zhou, J., Grimes, M. L., Bunnett, N. W., Howe, C. L. & Mobley, W. C. 1996. A signaling endosome hypothesis to explain NGF actions: potential implications for neurodegeneration. Cold Spring Harb Symp Quant Biol, 61, 389–406.

Bleasdale, J. E., Thakur, N. R., Gremban, R. S., Bundy, G. L., Fitzpatrick, F. A., Smith, R. J. & Bunting, S. 1990a. Selective inhibition of receptor-coupled phospholipase C-dependent processes in human platelets and polymorphonuclear neutrophils. J Pharmacol Exp Ther, 255, 756–68.

Bleasdale, J. E., Thakur, N. R., Gremban, R. S., Bundy, G. L., Fitzpatrick, F. A., Smith, R. J. & Bunting, S. 1990b. Selective-Inhibition of Receptor-Coupled Phospholipase-C-Dependent Processes in Human Platelets and Polymorphonuclear Neutrophils. Journal of Pharmacology and Experimental Therapeutics, 255, 756–768.

Bodmer, D., Ascano, M. & Kuruvilla, R. 2011. Isoform-Specific Dephosphorylation of Dynamin1 by Calcineurin Couples Neurotrophin Receptor Endocytosis to Axonal Growth. Neuron, 70, 1085–1099.

Bronfman, F. C., Lazo, O. M., Flores, C. & Escudero, C. A. 2014. Spatiotemporal intracellular dynamics of neurotrophin and its receptors. Implications for neurotrophin signaling and neuronal function. Handb Exp Pharmacol, 220, 33–65.

Cavalli, V., Kujala, P., Klumperman, J. & Goldstein, L. S. B. 2005. Sunday Driver links axonal transport to damage signaling. Journal of Cell Biology, 168, 775–787.

Cheng, Q., Song, S. H. & Augustine, G. J. 2017. Calcium-Dependent and Synapsin-Dependent Pathways for the Presynaptic Actions of BDNF. Frontiers in Cellular Neuroscience, 11.

Cheung, Z. H., Chin, W. H., Chen, Y., Ng, Y. P. & Ip, N. Y. 2007. Cdk5 Is Involved in BDNF-Stimulated Dendritic Growth in Hippocampal Neurons. PLoS Biol, 5, e63.

Cho, Y., Sloutsky, R., Naegle, K. M. & Cavalli, V. 2013. Injury-induced HDAC5 nuclear export is essential for axon regeneration. Cell, 155, 894–908.

Cohen, M. S., Bas Orth, C., Kim, H. J., Jeon, N. L. & Jaffrey, S. R. 2011. Neurotrophin-mediated dendrite-to-nucleus signaling revealed by microfluidic compartmentalization of dendrites. Proc Natl Acad Sci U S A, 108, 11246–51.

Cornejo, V. H., González, C., Campos, M., Vargas-Saturno, L., Juricic, M., Miserey-Lenkei, S., Pertusa, M., Madrid, R. & Couve, A. 2020. Non-conventional Axonal Organelles Control TRPM8 Ion Channel Trafficking and Peripheral Cold Sensing. Cell Rep, 30, 4505–4517.e5.

Cosker, K. E. & Segal, R. A. 2014. Neuronal signaling through endocytosis. Cold Spring Harb Perspect Biol, 6.

Delos Santos, R. C., Bautista, S., Lucarelli, S., Bone, L. N., Dayam, R. M., Abousawan, J., Botelho, R. J. & Antonescu, C. N. 2017. Selective regulation of clathrin-mediated epidermal growth factor receptor signaling and endocytosis by phospholipase C and calcium. Molecular Biology of the Cell, 28, 2802–2818.

Du, J., Feng, L. Y., Zaitsev, E., Je, H. S., Liu, X. W. & Lu, B. 2003. Regulation of TrkB receptor tyrosine kinase and its internalization by neuronal activity and Ca2+ influx. Journal of Cell Biology, 163, 385–395.

Du, J. L. & Poo, M. M. 2004. Rapid BDNF-induced retrograde synaptic modification in a developing retinotectal system. Nature, 429, 878–883.

Egan, M. F., Kojima, M., Callicott, J. H., Goldberg, T. E., Kolachana, B. S., Bertolino, A., Zaitsev, E., Gold, B., Goldman, D., Dean, M., Lu, B. & Weinberger, D. R. 2003. The BDNF val66met polymorphism affects activity-dependent secretion of BDNF and human memory and hippocampal function. Cell, 112, 257–69.

Gärtner, A., Polnau, D. G., Staiger, V., Sciarretta, C., Minichiello, L., Thoenen, H., Bonhoeffer, T. & Korte, M. 2006. Hippocampal long-term potentiation is supported by presynaptic and postsynaptic tyrosine receptor kinase B-mediated phospholipase Cgamma signaling. J Neurosci, 26, 3496–504.

Gonzalez, A., Moya-Alvarado, G., Gonzalez-Billaut, C. & Bronfman, F. C. 2016. Cellular and molecular mechanisms regulating neuronal growth by brain-derived neurotrophic factor. Cytoskeleton (Hoboken), 73, 612–628.

Goto-Silva, L., Mcshane, M. P., Salinas, S., Kalaidzidis, Y., Schiavo, G. & Zerial, M. 2019. Retrograde transport of Akt by a neuronal Rab5-APPL1 endosome. Sci Rep, 9, 2433.

Gruart, A., Sciarretta, C., Valenzuela-Harrington, M., Delgado-García, J. M. & Minichiello, L. 2007. Mutation at the TrkB PLC γ-docking site affects hippocampal LTP and associative learning in conscious mice. Learn Mem, 14, 54–62.

He, J., Gong, H. & Luo, Q. M. 2005. BDNF acutely modulates synaptic transmission and calcium signalling in developing cortical neurons. Cellular Physiology and Biochemistry, 16, 69–76.

Horch, H. W. & Katz, L. C. 2002. BDNF release from single cells elicits local dendritic growth in nearby neurons. Nat Neurosci, 5, 1177–84.

Huang, E. J. & Reichardt, L. F. 2003. Trk receptors: roles in neuronal signal transduction. Annu Rev Biochem, 72, 609–42.

Huang, S. H., Duan, S., Sun, T., Wang, J. E., Zhao, L., Geng, Z., Yan, J., Sun, H. J. & Chen, Z. Y. 2011. JIP3 Mediates TrkB Axonal Anterograde Transport and Enhances BDNF Signaling by Directly Bridging TrkB with Kinesin-1. Journal of Neuroscience, 31, 10602–10614.

Kang, H. & Schuman, E. M. 1995. Long-lasting neurotrophin-induced enhancement of synaptic transmission in the adult hippocampus. Science, 267, 1658–62.

Khamo, J. S., Krishnamurthy, V. V., Chen, Q., Diao, J. & Zhang, K. 2019. Optogenetic Delineation of Receptor Tyrosine Kinase Subcircuits in PC12 Cell Differentiation. Cell Chem Biol, 26, 400–410.e3.

Lang, S. B., Stein, V., Bonhoeffer, T. & Lohmann, C. 2007. Endogenous brain-derived neurotrophic factor triggers fast calcium transients at synapses in developing dendrites. Journal of Neuroscience, 27, 1097–1105.

Lazo, O. M., Gonzalez, A., Ascano, M., Kuruvilla, R., Couve, A. & Bronfman, F. C. 2013. BDNF regulates Rab11-mediated recycling endosome dynamics to induce dendritic branching. J Neurosci, 33, 6112–22.

Leal, G., Afonso, P. M., Salazar, I. L. & Duarte, C. B. 2015. Regulation of hippocampal synaptic plasticity by BDNF. Brain Research, 1621, 82–101.

Li, Y., Jia, Y. C., Cui, K., Li, N., Zheng, Z. Y., Wang, Y. Z. & Yuan, X. B. 2005. Essential role of TRPC channels in the guidance of nerve growth cones by brain-derived neurotrophic factor. Nature, 434, 894–898.

Lom, B., Cogen, J., Sanchez, A. L., Vu, T. & Cohen-Cory, S. 2002a. Local and target-derived brain-derived neurotrophic factor exert opposing effects on the dendritic arborization of retinal ganglion cells in vivo. J Neurosci, 22, 7639–49.

Lom, B., Cogen, J., Sanchez, A. L., Vu, T. & Cohen-Cory, S. 2002b. Local and target-derived brain-derived neurotrophic factor exert opposing effects on the dendritic arborization of retinal ganglion cells in vivo. Journal of Neuroscience, 22, 7639–7649.

Matsuda, N., Lu, H., Fukata, Y., Noritake, J., Gao, H., Mukherjee, S., Nemoto, T., Fukata, M. & Poo, M. M. 2009. Differential activity-dependent secretion of brain-derived neurotrophic factor from axon and dendrite. J Neurosci, 29, 14185–98.

Minichiello, L., Calella, A. M., Medina, D. L., Bonhoeffer, T., Klein, R. & Korte, M. 2002. Mechanism of TrkB-mediated hippocampal long-term potentiation. Neuron, 36, 121–37.

Moya-Alvarado, G., Guerra, M. V., Tiburcio, R., Bravo, E. & Bronfman, F. C. 2022. The Rab11-regulated endocytic pathway and BDNF/TrkB signaling: Roles in plasticity changes and neurodegenerative diseases. Neurobiol Dis, 171, 105796.

Moya-Alvarado, G., Tiburcio-Felix, R., Ibanez, M. R., Aguirre-Soto, A. A., Guerra, M. V., Wu, C., Mobley, W. C., Perlson, E. & Bronfman, F. C. 2023. BDNF/TrkB signaling endosomes in axons coordinate CREB/mTOR activation and protein synthesis in the cell body to induce dendritic growth in cortical neurons. Elife, 12.

Nguyen, M. K., Kim, C. Y., Kim, J. M., Park, B. O., Lee, S., Park, H. & Heo, W. D. 2016. Optogenetic oligomerization of Rab GTPases regulates intracellular membrane trafficking. Nat Chem Biol, 12, 431–6.

Redmond, L., Kashani, A. H. & Ghosh, A. 2002. Calcium regulation of dendritic growth via CaM kinase IV and CREB-mediated transcription. Neuron, 34, 999–1010.

Reichardt, L. F. 2006. Neurotrophin-regulated signalling pathways. Philos Trans R Soc Lond B Biol Sci, 361, 1545–64.

Rishal, I. & Fainzilber, M. 2014. Axon-soma communication in neuronal injury. Nature Reviews Neuroscience, 15, 32–42.

Scaramuzzino, C., Cuoc, E. C., Pla, P., Humbert, S. & Saudou, F. 2022. Calcineurin and huntingtin form a calcium-sensing machinery that directs neurotrophic signals to the nucleus. Sci Adv, 8, eabj8812.

Sciarretta, C., Fritzsch, B., Beisel, K., Rocha-Sanchez, S. M., Buniello, A., Horn, J. M. & Minichiello, L. 2010. PLC gamma-activated signalling is essential for TrkB mediated sensory neuron structural plasticity. Bmc Developmental Biology, 10.

Scott-Solomon, E. & Kuruvilla, R. 2018. Mechanisms of neurotrophin trafficking via Trk receptors. Mol Cell Neurosci, 91, 25–33.

Shelton, D. L., Sutherland, J., Gripp, J., Camerato, T., Armanini, M. P., Phillips, H. S., Carroll, K., Spencer, S. D. & Levinson, A. D. 1995. Human trks: molecular cloning, tissue distribution, and expression of extracellular domain immunoadhesins. J Neurosci, 15, 477–91.

Shimada, A., Mason, C. A. & Morrison, M. E. 1998. TrkB signaling modulates spine density and morphology independent of dendrite structure in cultured neonatal Purkinje cells. J Neurosci, 18, 8559–70.

Sholl, D. A. 1953. Dendritic organization in the neurons of the visual and motor cortices of the cat. J Anat, 87, 387–406.

Smallridge, R. C., Kiang, J. G., Gist, I. D., Fein, H. G. & Galloway, R. J. 1992. U-73122, an aminosteroid phospholipase C antagonist, noncompetitively inhibits thyrotropin-releasing hormone effects in GH3 rat pituitary cells. Endocrinology, 131, 1883–8.

Smith, R. J., Sam, L. M., Justen, J. M., Bundy, G. L., Bala, G. A. & Bleasdale, J. E. 1990. Receptor-coupled signal transduction in human polymorphonuclear neutrophils: effects of a novel inhibitor of phospholipase C-dependent processes on cell responsiveness. J Pharmacol Exp Ther, 253, 688–97.

Tapley, P., Lamballe, F. & Barbacid, M. 1992. K252a is a selective inhibitor of the tyrosine protein kinase activity of the trk family of oncogenes and neurotrophin receptors. Oncogene, 7, 371–81.

Taylor, A. M., Rhee, S. W., Tu, C. H., Cribbs, D. H., Cotman, C. W. & Jeon, N. L. 2003. Microfluidic Multicompartment Device for Neuroscience Research. Langmuir, 19, 1551–1556.

Xie, W., Zhang, K. & Cui, B. 2012. Functional characterization and axonal transport of quantum dot labeled BDNF. Integr Biol (Camb), 4, 953–60.

Xu, B., Zang, K., Ruff, N. L., Zhang, Y. A., Mcconnell, S. K., Stryker, M. P. & Reichardt, L. F. 2000. Cortical degeneration in the absence of neurotrophin signaling: dendritic retraction and neuronal loss after removal of the receptor TrkB. Neuron, 26, 233–45.

Zhang, Z. T., Fan, J., Ren, Y. X., Zhou, W. & Yin, G. Y. 2013. The release of glutamate from cortical neurons regulated by BDNF via the TrkB/Src/PLC-gamma 1 pathway. Journal of Cellular Biochemistry, 114, 144–151.

Zhou, B., Cai, Q., Xie, Y. & Sheng, Z. H. 2012. Snapin recruits dynein to BDNF-TrkB signaling endosomes for retrograde axonal transport and is essential for dendrite growth of cortical neurons. Cell Rep, 2, 42–51.

Zirrgiebel, U., Ohga, Y., Carter, B., Berninger, B., Inagaki, N., Thoenen, H. & Lindholm, D. 1995. Characterization of Trkb Receptor-Mediated Signaling Pathways in Rat Cerebellar Granule Neurons - Involvement of Protein-Kinase-C in Neuronal Survival. Journal of Neurochemistry, 65, 2241–2250.

